# Interleaved multi-magnification cryo-electron tomography bridges cellular and structural biology

**DOI:** 10.64898/2026.04.21.719848

**Authors:** Helena Watson, Victoria Garcia-Giner, Fabian Eisenstein, Michael Grange

**Affiliations:** The Rosalind Franklin Institute, Harwell Science & Innovation Campus, Didcot OX11 0QS, United Kingdom; Institute of Molecular Biology and Biophysics, ETH Zürich, Zürich, Switzerland; Division of Structural Biology, Wellcome Centre for Human Genetics, University of Oxford, OX3 7BN Oxford, United Kingdom; DECTRIS Ltd., Taefernweg 1, 5405 Baden-Daettwil, Switzerland

## Abstract

Cryo-electron tomography (cryo-ET) enables in situ structural analysis of macromolecular complexes within native cellular environments. However, the limited field of view required for high-resolution structure determination necessarily restricts a wider assessment of the broader cellular context. We present a multi-magnification cryo-ET acquisition strategy that integrates low- and high-magnification information from coincident sample regions during the same tilt-series. By interleaving acquisition of the magnifications at each tilt angle, this strategy enables simultaneous collection of large field-of-view, low-resolution tomograms and high-resolution, small field-of-view tomograms while minimising the impact of the increased electron dose. We demonstrate that we can capture cellular organisation across tens of microns, while still enabling subtomogram averaging to resolutions below 4 Å. This integrated acquisition framework establishes a practical route to multi-scale cryo-ET, bridging molecular and cellular scales for more comprehensive biological insight.

## Introduction

A holistic understanding of the molecular organisation within a biological system requires integration of structural information across multiple spatial scales, from atomic-level molecular interactions to the organisation of cells and tissues. Biological processes arise from coordinated interactions of components across these scales, making it essential to relate molecular architecture to cellular and multicellular context.

Cryo-electron tomography (cryo-ET) enables the high-resolution study of natively preserved, frozen-hydrated biological samples, allowing macromolecular structures to be visualised directly within their native cellular environment. Recent advances in focused ion beam (FIB) milling have further expanded the scope of cryo-ET, enabling access to increasingly complex specimens, including multicellular, tissue and organism samples (Schiøtz et al., 2024; Nguyen et al., 2024; Glynn et al., 2025). As the study of such biologically complex systems by cryo-ET becomes increasingly routine, it is important to integrate high-resolution molecular information with broader ultrastructural context to maximise insight from these samples.

The preparation of lamellae by cryo-FIB milling exposes electron-transparent surface areas of thousands of square microns, orders of magnitude larger than the field of view typically captured by cryo-ET (Berger et al., 2025; Glynn et al., 2025). The high magnifications required for high-resolution structure determination constrain the two-dimensional field of view of each tilt-series, often to less than 1 µm^2^. Although parallelised beam-shift data acquisition strategies have increased throughput (Bouvette et al., 2021; Eisenstein et al., 2022), the limited field of view still restricts the overall imaging throughput across lamellae. Consequently, much of the biological information within a lamella remains unsampled. While such high magnifications and resolutions are not always necessary to address a given biological question, current cryo-ET workflows effectively require a compromise at the point of data acquisition between resolution and spatial coverage, restricting the ability to effectively interpret molecular structures within their native cellular environment.

A further consideration is the radiation sensitivity of frozen-hydrated samples. Cumulative electron dose during tilt-series acquisition leads to progressive loss of high-resolution information and sample deformation (Leapman and Sun, 1995; Grant and Grigorieff, 2015). Acquisition schemes must therefore be carefully designed to preserve high-resolution signal (Hagen et al., 2017), placing strict constraints on how additional contextual information can be obtained. Although approaches such as montage tomography have been developed to increase effective FOV by tiling high-magnification images (Peck et al., 2022; Yang et al., 2023), overlapping exposures lead to increased dose accumulation and can limit the resolution attainable by subtomogram averaging (Hylton et al., 2025). In addition, montage strategies substantially increase data volume and processing complexity.

We present a **B**road **I**nformation **G**athering **S**trategy by **M**ultiscale **A**cquisition of lame**LL**a (BIGSMALL). This enables parallel, interleaved collection of high- and low-magnification tilt-series from the same region of interest. Rather than expanding the field of view through high-magnification tiling, this approach directly captures large-scale cellular context using low-magnification data while simultaneously acquiring targeted high-resolution datasets suitable for subtomogram averaging. This strategy provides a direct route to spatially contextualise high-resolution structural information within larger cellular volumes, without relying on separate imaging modalities or compromising high-resolution data quality. By bridging molecular detail and cellular organisation within a multi-scale acquisition framework, this approach advances cryo-ET to enable a more integrated understanding of biological systems.

## Results

### BIGSMALL enables integrated multi-scale cryo-ET across molecular and cellular scales

To improve the contextualisation of high-resolution cryo-ET data, we developed BIGSMALL, a multi-magnification acquisition strategy in which low-magnification (LM) and high-magnification (HM) tilt-series are acquired from coincident regions within thinned lamellae (Figure 1A).

**Figure 1.**
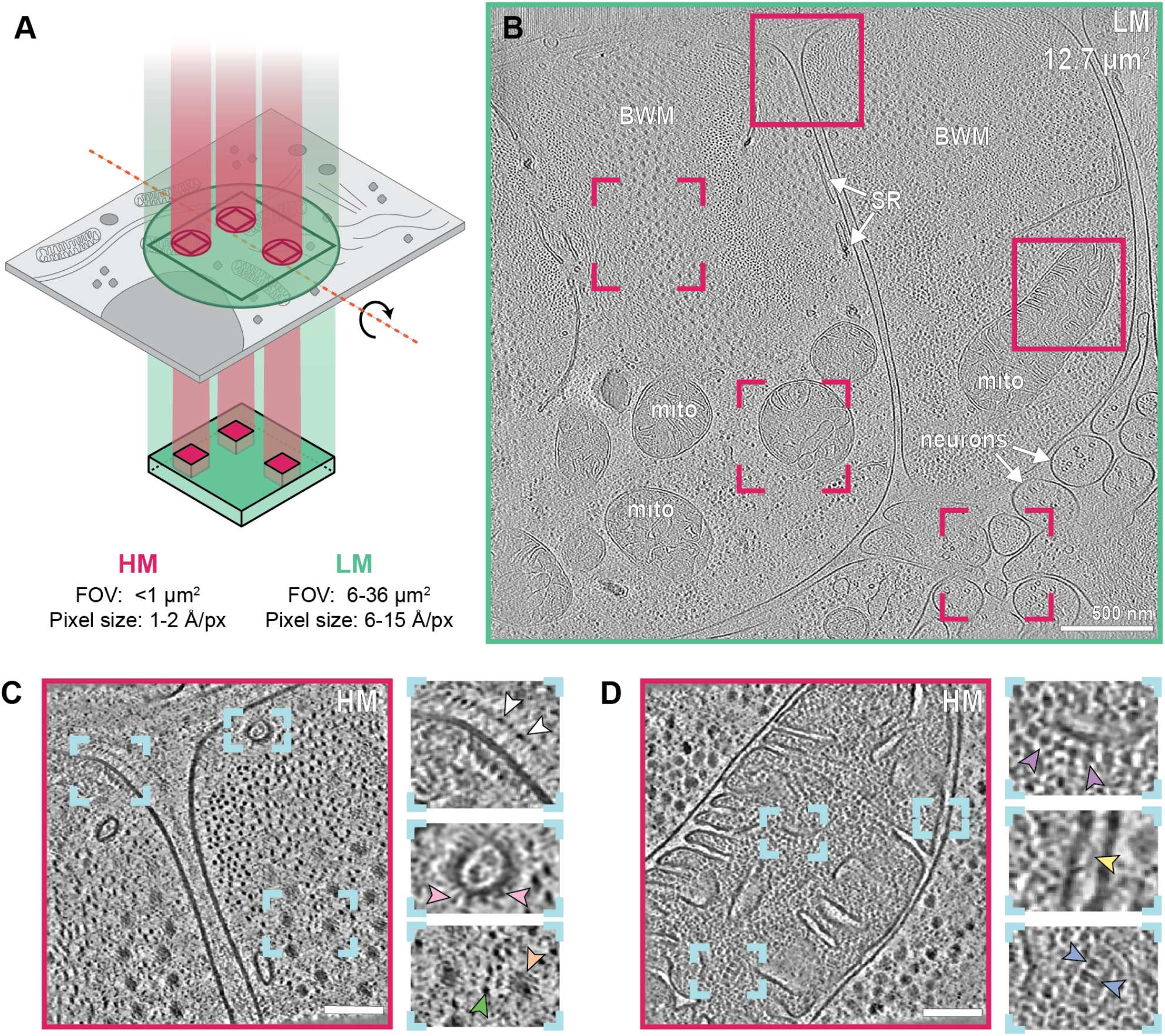
Multi-scale cryo-ET enables integrated cellular and molecular imaging. **(A)**Schematic of multi-magnification tilt-series acquisition. At each tilt, low-magnification (green) and high-magnification (magenta) images are acquired from coincident sample areas. Independent tomographic reconstructions yield large field-of-view, low resolution tomograms and small field-of-view, high-resolution tomograms. **(B)**Low-magnification tomogram slice of *C. elegans* body wall muscle cells acquired with BIGSMALL (0.3 e^—^ /Å^2^/tilt), with regions targeted for HM tomograms outlinedin magenta. Examples of body wall muscle cells (BWM), neurons, mitochondria (mito) and sarcoplasmic reticulum (SR) are annotated. Scale bar: 500 nm. (**C**,**D**) Example denoised high-magnification tomogram slices from regions shown with solid boxes in (B), with insets showing magnified views of areas marked in blue. Scale bars: 100 nm. Arrowheads indicate subcellular features: white, attachment plaque protein; pink, SERCA proteins; orange, thick muscle filaments; green, thin muscle filaments; purple, ATP synthase; yellow, prohibitin; blue, stacked mitochondrial cristae.

High-magnification cryo-ET acquisition provides high-resolution structural information, with small pixel sizes (typically <2 Å per pixel) enabling subtomogram averaging to resolutions beyond 4 Å (Xing et al., 2023; Xue et al., 2025). In contrast, low-magnification acquisition uses larger pixel sizes, yielding lower-resolution data but a substantially expanded field of view that captures the wider cellular architecture within the lamella. As a result, multiple high-magnification tilt series can be acquired within the field of view of a single low-magnification tilt series, allowing efficient and spatially correlated multi-scale sampling (Figure 1B).

We demonstrate this strategy using lamellae prepared from adult *C. elegans*, where five high-magnification tilt series were acquired within the field of view of a single low-magnification tilt series. The low-magnification tomogram provides a broad overview of tissue organisation (Figure 1B), allowing identification and assignment of cellular features based on the well-defined anatomy of *C. elegans*, such as body wall muscle cells and adjacent neurons. At this scale, global sarcomere organisation is readily observed, and organelles such as mitochondria and sarcoplasmic reticulum can be readily identified across the tomogram.

High-magnification tomograms provide detailed structural information from spatially constrained regions of interest (Figure 1C,D; Supplementary Figure S1). For example, when focusing on the interfaces between adjacent muscle cells, membrane-associated densities are resolved, including finger-like structures at putative attachment plaque sites (Qadota et al., 2017) and complexes consistent with SERCA pumps on the sarcoplasmic reticulum (Chen and Kudryashev, 2020) (Figure 1C). In mitochondria, high-magnification tomograms reveal inner mitochondrial membrane complexes, including ATP synthase (Strauss et al., 2008) and prohibitin assemblies (Lange et al., 2025) (Figure 1D). The small pixel size of HM data enables direct identification of these complexes and makes the data directly amenable to subtomogram averaging. Critically, the LM tomogram provides the surrounding cellular information that is not captured within the limited HM field of view. For instance, while a HM tomogram may only encompass a portion of a mitochondrion, the corresponding LM dataset reveals its full extent and spatial relationship to neighbouring cellular structures (Figure 1B,D).

Low-magnification tomograms acquired with BIGSMALL also enable direct three-dimensional analysis of cellular architecture at scales not accessible by high-magnification acquisition. In *C. elegans* lamellae, LM tomograms reveal that the rough endoplasmic reticulum (RER) is highly organised into extended stacks of cisternae, which in some regions are arranged in concentric layers (Figure 2A ; Supplementary Figure S2). Three-dimensional segmentation shows that the RER network is highly fenestrated, containing numerous pores and localised membrane constrictions across the cisternae (Figure 2B,C). These features are consistent with the established view of the ER as a continuous and highly dynamic membrane network, capable of adopting diverse morphologies depending on cellular context (Friedman and Voeltz, 2011). Importantly, these fenestrations can be mapped across extended regions of the ER, enabling mesoscale structural organisation to be analysed over micron-scale volumes (Figure 2D). Previous studies have demonstrated that ER architecture varies with subcellular location and cell cycle state, including transitions between sheet-like and tubular morphologies and changes in the degree of fenestration (Puhka et al., 2012; Shemesh et al., 2014). Thus, cellular architecture at this scale reflects the biological state of the specific cell within the organism at the time of vitrification. The ability to characterise these features in 3D over larger volumes provides a route to correlate ER ultrastructure with local cellular context, including proximity to other organelles or subcellular domains. Crucially, this analysis relies on volumetric data. Such spatial relationships cannot be resolved using two-dimensional projection images of lamellae, which lack depth information and obscure membrane connectivity. Instead, low-magnification tilt-series acquisition and three-dimensional reconstruction are required to capture these mesoscale features.

**Figure 2.**
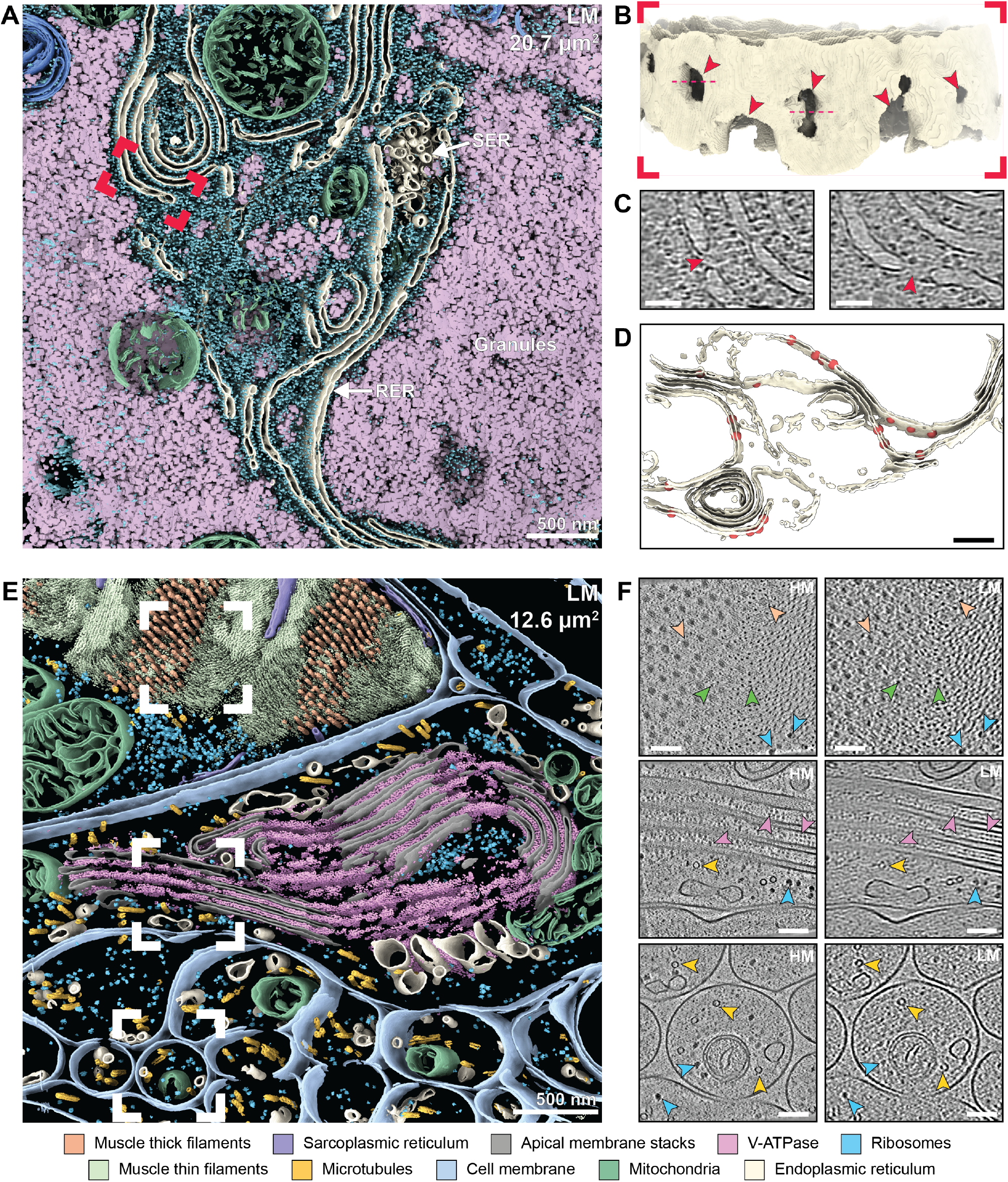
Low-magnification cryo-ET resolves cellular architecture with molecular continuity to high magnification. (**A**) Segmentation of an LM tomogram highlighting RER and other organelles. Scale bar: 500 nm. (**B**) Magnified view of RER in the region indicated by the red inset in (A). Red arrowheads mark pores in the RER membrane. (**C**) LM tomogram slices from regions indicated by dotted lines in (B), with red arrowheads highlighting the same RER membrane pores. Scale bar: 100 nm. (**D**) Back-projection of pore positions, represented by spheres, onto the RER membrane segmentation. Scale bar: 500 nm. (**E**) Segmentation of an LM tomogram of *C. elegans* tissue, show muscle cells, hypodermis, neurons. Scale bar: 500 nm. (**F**) Comparison of features visible in HM and LM tomogram slices (cropped to same FOV). Arrowheads indicate macromolecular features visible at both magnifications: orange, thick muscle filaments; green, thin filaments; blue, ribosomes; pink, ATPases; yellow, microtubules. Scale bar: 100 nm.

BIGSMALL integrates the distinct and complementary strengths of low- and high-magnification within a single experimental framework. By using intermediate pixel sizes (6-15 Å/px), low-magnification tomograms capture fields of view spanning tens of µm^2^ (Figure 2E) while still retaining sufficient resolution to visualise individual macromolecules (Figure 2F). Notably, many large macromolecules are visible at both magnifications, providing a direct bridge between scales. Structures such as muscle filaments, ribosomes, ATPases and microtubules can be identified in LM tomogram slices and resolved in greater detail in corresponding HM tomogram slices (Figure 2F). As LM and HM datasets are acquired from coincident regions of a lamella, the datasets are intrinsically co-localised, allowing structural information to be interpreted across scales at precisely defined spatial positions within an extended cellular landscape.

### A dose efficient interleaved acquisition strategy for multi-scale cryo-ET

A key consideration in multi-scale cryo-electron tomography is the impact on vitrified biological specimens of added exposure to the electron beam. However, while high-magnification (HM) imaging typically requires 3-5 e^—^/Å^2^/tilt, low-magnification imaging can be performed at substantially lower doses (0.2-0.3 e^—^/Å^2^/tilt), aided by increased defocus to enhance low-frequency contrast (Supplementary Figure S3, Supplementary Table S1). As a result, a complete LM tilt series can be acquired with a cumulative dose below 15 e^—^/Å^2^. This corresponds to less than 10% of that required for a conventional HM tilt-series. Therefore in principle, this enables combined multi-scale acquisition with only a modest increase in total dose.

A further consideration is acquisition order of multiscale tilt series, as sequential acquisition of LM and HM datasets can introduce significant costs to data quality. For example, the dose accumulated by acquiring HM data first (>120 e^—^/Å^2^) can introduce large-scale sample deformations or hydrogen bubble formation (Leapman and Sun, 1995), compromising interpretation of subsequent LM tomograms. Conversely, the dose corresponding to acquiring LM data first (<15 e^—^/Å^2^) leads to degradation of high-resolution information in the HM dataset (Grant and Grigorieff, 2015), limiting the resolution at which structures may be determined by subtomogram averaging.

To address this, we implemented an interleaved acquisition strategy, in which LM and HM images are collected at each tilt angle throughout the tilt series (Figure 3A). Dose-symmetric acquisition is now widely adopted as the most dose-efficient method for subtomogram averaging (Hagen et al., 2017), and forms the basis for our interleaved acquisition strategy. Our approach distributes the total dose symmetrically across both low and high magnification tilt series and ensures that the minimal additional dose is applied at low tilt angles, where high-resolution information is most critical (Figure 3B). Consequently, the impact of prior electron exposure on both high-resolution structural information and larger-scale cellular context is minimised. Importantly, interleaving ensures that both datasets capture the specimen in an identical structural state at each tilt angle, enabling direct and reliable correlation across spatial scales. Additionally, this approach preserves acquisition throughput, minimising stage movement to a single tilt for each LM/HM pairing.

**Figure 3.**
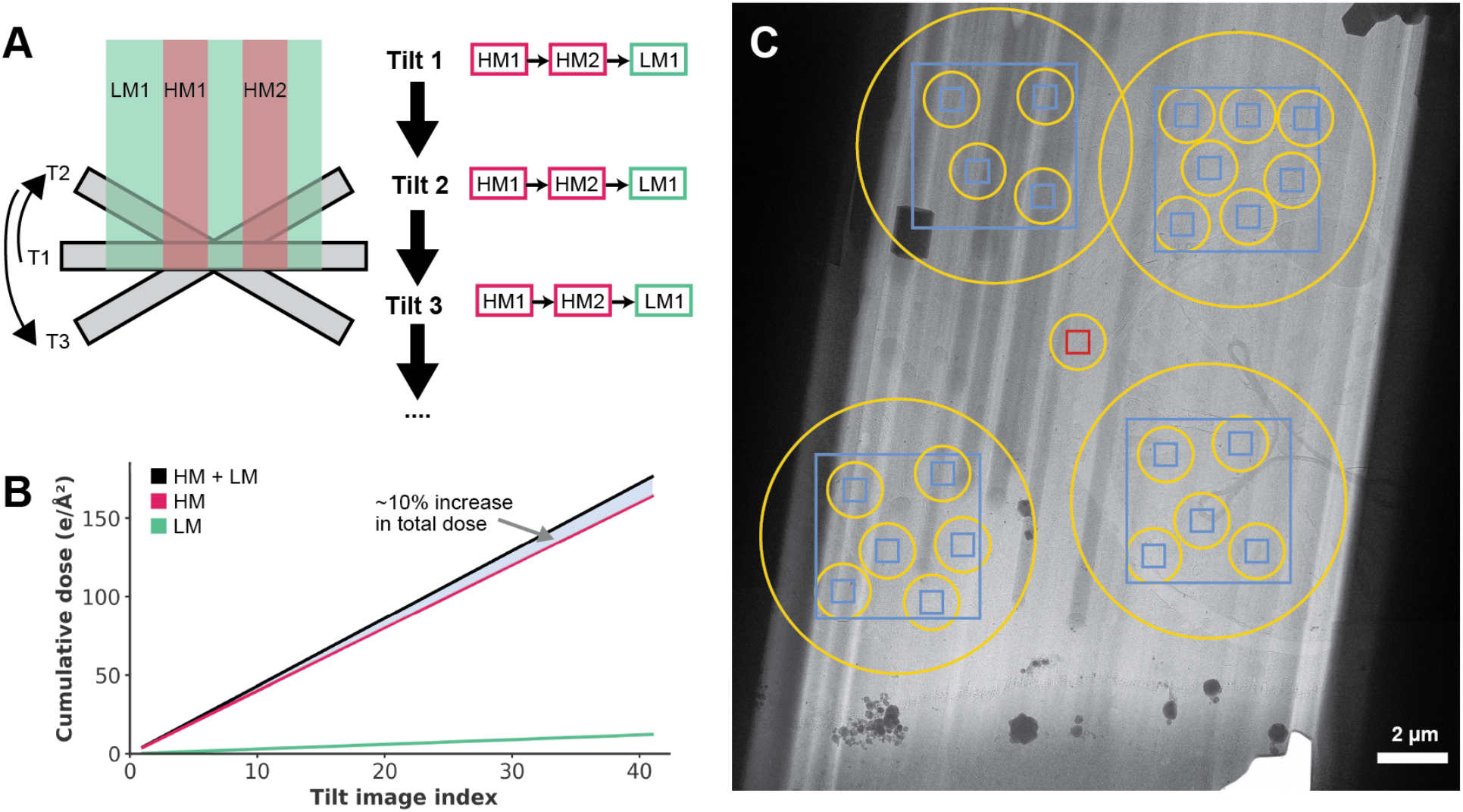
Implementation of interleaved multi-magnification cryo-ET. (**A**) Schematic of the interleaved acquisition strategy. At each stage tilt, images at all magnifications are collected before proceeding to the next stage tilt angle. High-magnification (HM, pink) and low-magnification (LM, green) images are acquire sequentially within each tilt. (**B**) Cumulative electron dose as a function of tilt image index. The additional dose from LM imaging is minimal, typically contributing <10% to the total accumulated dose. (**C**) Example of the targeting setup. Blue squares indicate acquisition areas, the red square marks the high-magnification tracking and focus area, and yellow circles denote the illuminated regions for each acquisition.

BIGSMALL was implemented within PACEtomo (Eisenstein et al., 2022), an automated parallel beam-shift tilt-series acquisition framework built on the open-source microscope control software SerialEM (Mastronarde, 2005; Schorb et al., 2019). The approach extends existing workflows to enable multi-magnification acquisition by allowing targets at different magnifications to be defined within a single acquisition group (Figure 3C; Supplementary Figure S4). During acquisition, all targets within a group are imaged sequentially at each tilt angle according to their assigned imaging state using a beam shift. This enables interleaved multi-magnification acquisition without additional stage movements and preserves standard dose-symmetric tilt schemes. The approach therefore retains the efficiency of conventional parallel beam-shift acquisition workflows. The framework is readily extensible to additional magnifications, provided that cumulative dose constraints are considered. More generally, this implementation requires only the ability to assign distinct imaging states to targets within a shared acquisition sequence and is therefore transferable to other automated beam-shift acquisition software. As an alternative approach to interleaved acquisition (Downes et al., 2025), the same framework also supports non-interleaved acquisition by defining separate target groups for different magnifications and ordering the target groups appropriately.

### Interleaved acquisition preserves high-resolution in situ subtomogram averaging

The dose required for low-magnification imaging is inherently small, and the interleaved acquisition strategy is designed to minimise its impact on high-resolution information while preserving acquisition throughput. To directly assess whether interleaving LM imaging affects the attainable resolution of subtomogram averaging, we collected in situ datasets from *E. coli* lamellae under matched conditions. Conventional high-magnification tilt series (4 e^—^/Å^2^/tilt) were compared to high-magnification tilt series acquired interleaved with low-magnification imaging at 0.5 e^—^/Å^2^/tilt. This LM dose lies at the upper end of typical values (more commonly 0.2-0.3 e^—^/Å^2^/tilt), representing a conservative test of the impact of additional exposure and approximating scenarios in which multiple LM exposures may overlap.

Subtomogram averaging of ribosomes from HM tomograms was performed independently for both datasets. To minimise the impact of conformational heterogeneity arising from different translational states, refinement was focused on the 50S large ribosomal subunit. Both acquisition strategies yielded reconstructions at a global 50S resolution of 3.7 Å, with extensive regions of the 50S subunit reaching local Nyquist-limited resolutions of 3.0 Å (Figure 4A,B; Supplementary Figure S6). Visual inspection of the resulting maps revealed comparable preservation of high-resolution features. In particular, side-chain densities were clearly resolved in both reconstructions, including densities corresponding to radiation-sensitive acidic residues (Figure 4C).

**Figure 4.**
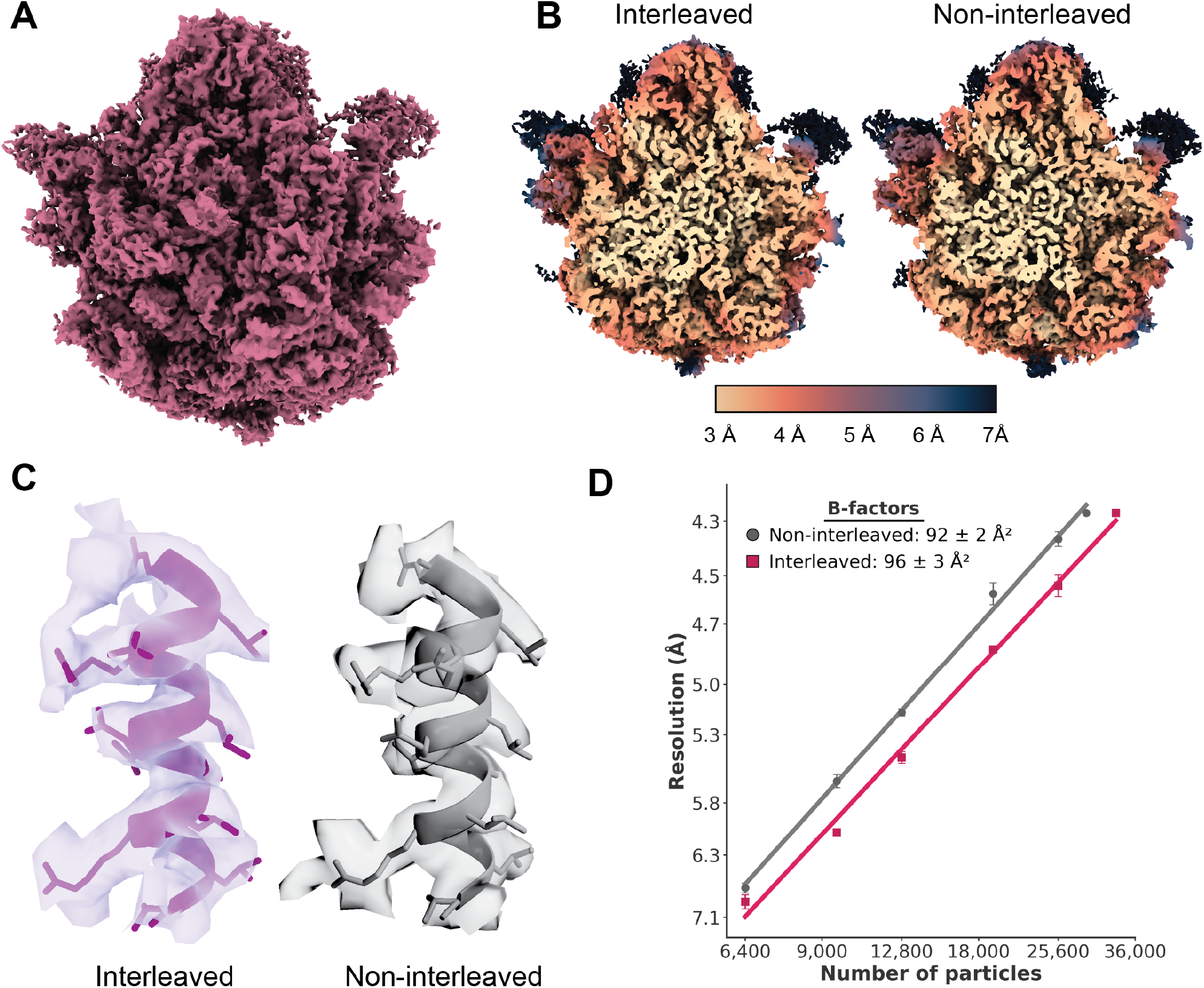
Interleaved acquisition is compatible with high-resolution structure determination. **(A)** Density map of the *E. coli* 50S ribosomal subunit average from interleaved (BIGSMALL) acquisition. **(B)** Central slices through interleaved and non-interleaved density maps, coloured by local resolution. **(C)** Representative rigid model fit (ribosomal L13 chain, PDB: 6GC8 (Nikolay et al., 2018)) into the density maps of interleaved and non-interleaved reconstructions. **(D)** B-factor analysis of interleaved and non-interleaved datasets. Resolution (1/R^2^ scale, mean of n = 3 technical replicates) is plotted against particle number (log_e_scale).

Consistent with this, comparison of the B-factors to control for differences in particle number (Rosenthal and Henderson, 2003) derived from the two reconstructions revealed only minor differences, with values of 96 Å^2^ for the interleaved dataset and 92 Å^2^ for the non-interleaved dataset (Figure 4D). As both datasets were acquired from the same lamellae and exhibited similar thickness distributions (Supplementary Figure S7), the small differences in B-factor indicate that the additional dose associated with interleaved LM acquisition results in minimal dampening of high-resolution signal. Together, these results demonstrate that interleaved multi-magnification acquisition is compatible with high-resolution in situ structure determination by STA.

## Discussion

BIGSMALL enables the direct integration of molecular and cellular information by interleaving low- and high-magnification cryo-electron tomography within a single acquisition. In this approach, high-resolution tomograms are embedded within larger contextual volumes, establishing continuity across length scales and preserving spatial relationships from nanometre-scale molecular structures to micron-scale cellular architecture.

This strategy bridges cellular and structural biology within a unified framework. Low-magnification tomograms provide three-dimensional ultrastructural information over micron-scale areas, capturing features such as membrane topology, organelle organisation and tissue architecture. For example, resolving the connectivity of endoplasmic reticulum networks, including fenestrations and stacked sheet organisation, requires a field of view that extends beyond that accessible at high magnification (Figure 2). High-magnification tomograms resolve macromolecular complexes and their structural states, such as ribosomal interactions. By integrating these complementary data, BIGSMALL enables structural heterogeneity to be interpreted while incorporating its subcellular localisation, organisation, and heterogeneous cellular state. This establishes a framework for spatially resolved analysis of macro-molecular organisation and for relating molecular-scale perturbations to higher-order cellular and tissue architecture, which cannot be achieved through individual imaging regimes alone.

Conceptually, BIGSMALL shares similar goals with montage tomography approaches, which also aim to increase the field of view. However, montage strategies incur significant practical challenges. Covering large areas at high magnification requires the acquisition of many overlapping tilt series, leading to substantial cumulative electron dose and increased radiation damage in localised regions (Hylton et al., 2025). Mitigating these effects often necessitates specialised hardware modifications (Chua et al., 2024) or complex acquisition schemes (Peck et al., 2022; Yang et al., 2023), with the size of the resultant datasets posing significant issues for data processing. In contrast, BIGSMALL provides a more efficient solution by leveraging the dose efficiency of low magnification imaging and interleaving acquisitions. This strategy requires a substantially reduced number of tilt series to sample a given region, enabling larger fractions of the lamella to be explored without prohibitive increases in dose or data volume. BIGSMALL therefore increases effective sampling of biological material, which is particularly advantageous for heterogeneous or low-throughput specimens such as tissues or multicellular systems, where maximising information yield from each lamella is critical.

Developments in instrumentation are likely to further expand the capabilities of BIGSMALL. Advances in chromatic aberration correction (Dickerson et al., 2022; Wu et al., 2025) and laser phase plate technologies (Schwartz et al., 2019; Du and Fitzpatrick, 2023; Petrov et al., 2024) are expected to enable imaging of thicker specimens, extending the accessible volume in the axial dimension. These combined developments will expand the ability to capture larger, more representative biological volumes within a single experiment.

Realising the full potential of multi-scale cryo-ET will require robust approaches for integrating and analysing data across magnifications. Standardisation of coordinate systems, metadata, and transformation parameters will be essential to maintain spatial relationships between datasets and enable reproducible analysis. This need extends beyond multi-scale cryo-ET to include correlative imaging workflows, such as CLEM, as well as lamella overview maps that provide critical spatial context. We anticipate that data repositories and analysis platforms will increasingly need to support explicitly linked multi-scale datasets, in which annotated regions of interest can be analysed across datasets to enable comparative analysis.

Together, BIGSMALL establishes multi-scale cryo-ET as a practical framework for structural cell biology, enabling integrated analysis of molecular architecture and cellular organisation within native environments. By combining complementary information across imaging scales and increasing the effective sampling of biological specimens, this approach provides a route towards a more comprehensive and representative understanding of how molecular assemblies give rise to cellular function.

## Methods

### *E. coli* sample preparation

*E. coli* C43 (DE3) were incubated in LB at 37 °C in a shaking incubator and grown to an OD_600_ of 0.6 to recover exponential growth. The bacteria were then re-suspended with glycerol in LB to a 10% (*v/v*) final concentration of glycerol and concentrated by centrifugation at 1000 × *g* immediately before vitrification.

Bacteria were vitrified by high-pressure freezing in a Leica EM ICE (Leica Microsystems). 6 mm planchettes (Au-plated Cu carrier type B, Leica) were polished with fine grit sandpaper up to a final grit of 10,000 and metal polish before coating the flat sides of the planchettes with 1% (*w/v*) soy lecithin solution in chloroform. Samples were assembled for high pressure freezing with the following geometry (bottom to top): planchette (flat side up), 200 mesh Cu grid (Agar Scientific) glow-discharged on each side for 45 s at 30 mA, 10-15 µl bacterial pellet, planchette (flat side down). Samples were disassembled under liquid nitrogen and the vitrified grids were clipped into Autogrids (Thermo Fisher Scientific).

### *E. coli* lamella preparation

Focused ion beam milling was performed with an Arctis PFIB/SEM (Thermo Fisher Scientific) using xenon plasma at 30 kV as previously described (Berger et al., 2025). Ice contamination was removed from both sides of the grid by FIB imaging at 15 nA at the lowest possible magnification. After ice removal, the front side of the grid was sputter-coated with platinum for 120 s at 12 kV and 70 nA, followed by deposition of organometallic platinum for 120 s using the gas injection system, then another platinum sputter-coating step for 120 s.

Trench milling was performed from the back side of the grid, perpendicular to the plane of the grid. Trenches measuring 25 × 30 µm were milled using cleaning cross-section (CCS) mode at 60 nA, with a distance of 25 µm separating each pair of trenches. From the front side of the grid, material below the intended lamella site was removed at stage tilts of 45°, 28°and 20°relative to the grid plane using a beam current of 15 nA.

Lamella sites were identified in Maps 3.21 (Thermo Fisher Scientific) and imported into AutoTEM Cryo 2.4 (Thermo Fisher Scientific) for automated lamella preparation with a target milling angle of 20° relative to the grid plane. Milling was performed at progressively decreasing beam currents of 4.0 nA, 1.0 nA and 0.3 nA, with a corresponding reduction in the distance between the milling pattern and the intended lamella site. Over- and under-tilts of ±1° and ±0.5° were applied during the 1.0 nA and 0.3 nA milling steps respectively. Final lamella polishing was performed at 10 pA.

### *E. coli* tilt-series acquisition

Tilt-series were acquired on a Titan Krios G4 transmission electron microscope (Thermo Fisher Scientific) equipped with a Selectris X energy filter. Data were recorded on a Falcon 4i camera in EER format using electron counting mode. Data collection was performed in SerialEM (v4.2.1) using SPACEtomo (v1.3) (Eisenstein et al., 2024), with beam tilt compensation (using SerialEM’s Coma vs Image Shift calibration). Tilt series were collected using a dose-symmetric scheme with a starting tilt of -20°to correct for the lamella milling angle, with a tilt range of ±54°with 3°increments.

High-magnification tilt series were acquired a nominal magnification of 81,000x (calibrated pixel size 1.48 Å/px) using the SerialEM Low Dose Record preset. The electron dose was 4 e^—^/Å^2^/tilt image, with a target defocus range of -3 to -5 µm applied in 0.5 µm increments. Low-magnification tilt series were acquired at 15,000x magnification (pixel size 8.42 Å/px) using the SerialEM Low Dose Search preset, with an electron dose of 0.5 e^—^/Å^2^/tilt image and a defocus offset of -20 µm relative to the high-magnification setting.

Two experimental conditions were collected for comparison. In the non-interleaved condition, target groups contained only high-magnification (HM; SerialEM Record) targets. In the interleaved condition, target groups contained both HM (Record) and low-magnification (LM; SerialEM Search) targets, such that HM and LM images were acquired at each stage tilt. To ensure that LM imaging did not affect the HM-only dataset, all HM-only tilt series for a given lamella were acquired prior to the interleaved acquisitions for that lamella. For interleaved datasets, HM targets were positioned within the illumination area of the corresponding LM target.

### *E. coli* cryo-ET data processing

Gain correction, motion correction and CTF estimation of tilt images was performed using the WarpTools pipeline (Tegunov and Cramer, 2019). Tilt images in which grid bars or significant ice contamination entered the field of view were manually excluded using custom scripts. Tilt-series alignment and tomogram reconstruction was performed using AreTomo (v1.2.5) (Zheng et al., 2022). High-magnification datasets were reconstructed with a binning factor of 8 (11.84 Å/px). Reconstructed tomograms were flipped using IMOD (Mastronarde and Held, 2017) to obtain the correct map handedness, and subsequently either low-pass filtered to 60 Å or deconvolved with a Wiener-like filter using IsoNet1 (Liu et al., 2022), employing a model previously trained on *E. coli* tomograms (Berger et al., 2025).

A crYOLO model (Wagner et al., 2019) was trained on ribosome particles manually picked from a deconvolved tomogram and subsequently applied to all deconvolved tomograms to generate initial particle coordinates. Particles derived from tomograms acquired in interleaved and non-interleaved modes were processed independently following the same workflow. Subvolumes were reconstructed using WarpTools at a binning factor of 4 (pixel size 5.92 Å/px) with a cubic box size of 80^3^ voxels. Subtomograms were refined in RELION (v4) using a non-aligned average as the initial reference. Following 3D classification, selected particles were subjected to further 3D refinement. Particles were then re-extracted in WarpTools at a binning factor of 2 (pixel size 2.96 Å/px) and refined further in RELION. Further refinement of particles was performed in M (Tegunov et al., 2021), including iterative cycles of particle pose refinement, image warp refinement (up to 4!4), CTF refinement, volume warp refinement (4!4!2!10), stage angle refinement, and particle temporal trajectory refinement. Per-tilt-series weighting and LSU-focused masking were applied, resulting in both datasets converging to a final resolution of 3.7 Å for the LSU, as determined by the gold-standard FSC 0.143 criterion.

B-factor analysis for both datasets was performed in RELION. Subtomograms were reconstructed using WarpTools at a binning factor of 1 (1.48 Å/px) with a box size of 280^3^ voxels. Random particle subsets were generated and refined using the RELION script relion_bfactor_plot.py, with a minimum particle number of 6,400. For each dataset, the B-factor analysis was repeated three times, yielding n = 3 resolution measurements for each particle subset. The B-factor was calculated as 2 divided by the slope of the linear fit between log_e_(number of particles) and 1/resolution^2^.

### *C. elegans* sample preparation

Wild-type and mutant (*dnj-25 (R743G)*) *C. elegans* strains expressing a dopaminergic neuron GFP reporter (*dat-1p::GFP*) were used in this study. Strains were provided by the Kevei Group at the University of Reading. Nematodes were cultivated on NGM plates seeded with *E. coli* OP50 according to standard protocols. Populations were synchronised using the bleaching method; the resulting egg solution was washed with M9 buffer and incubated at 20 °C for 24 hours to allow the eggs to hatch. The L1 larvae were then seeded onto 6 cm NGM plates at a density of 200-300 larvae per plate and grown until the day 1 adult stage (60-72 hrs).

Adult worms from a 6 cm plate were resuspended in 2 ml M9 buffer and allowed to settle by gravity (<5 min). The supernatant was removed, leaving 50 µL of worm pellet, which was resuspended with cryoprotectant to a final concentration of 10% (*w/v*) dextran, 10% (*w/v*) trehalose and 5 mM levamisole in M9. After 5-10 mins incubation, 3 µl of *C. elegans* suspension was applied to the 100 µm cavity of a 3 mm Type A Au-coated Cu planchette (Wohlwend). A polished Type B planchette (flat side coated with 1-hexadecene) was then placed flat side down to close the assembly. The assembled carriers were immediately vitrified by high-pressure freezing in a Leica EM ICE.

Vitrified sample carriers were screened on a Stellaris 8 Cryo Confocal Microscope (Leica Microsystems, HC PL APO 50 x0.9-NA objective) to assess ice quality and locate worms of interest using widefield GFP fluorescence and reflection imaging. Tilesets were acquired using LASX software (Leica Microsystems).

### *C. elegans* lamella preparation

Samples were loaded onto a 27°shuttle and transferred to a Helios G5 Hydra CX plasma FIB/SEM (Thermo Fisher Scientific). Samples were oriented according to cryo-fluorescence screening to facilitate the lift-out of the nematode of interest. Planar serial lift-out was performed using xenon plasma at 30 kV (Glynn et al., 2025). Fiducial marks were milled at 4 nA to facilitate navigation, and *C. elegans* were identified via fluorescence imaging with the integrated Delmic METEOR system (100 mW, 100 ms). The sample was coated sequentially with sputtered platinum (60 s), organometallic platinum delivered via the gas injection system (60 s), and a further layer of sputtered platinum (60 s). Subsequent fluorescence imaging was performed at 300 mW and 300 ms. With the sample oriented perpendicular to the FIB, a front trench of 100 × 300 µm was milled with a rectangular cross-section (RCS) pattern using currents between 0.2 µA and 60 nA. Side trenches (7 × 170 × 20 µm) were milled at 60 nA, followed by an undercut (70 × 3 × 50 µm) milled parallel to the FIB at the shallowest possible angle (6° to 8°) using a 15 nA beam. The sample was then rotated to be perpendicular to the FIB again, and the sample attached to the copper block adaptor on the EasyLift needle with 5-6 RCS patterns of 0.8 × 2.5 × 5 µm with 4 µm periodicity, milled at 0.3 nA. The extraction volume was then released by milling the last connection to the bulk with rectangular patterns at 15 nA, and the EasyLift needle with attached sample was retracted. Serial sections were deposited at a glancing angle onto a 400 × 100 mesh copper TEM support grid (Agar Scientific), with individual chunk dimensions of 42µm width and 4µm height. Following deposition, the stage was tilted to steeper milling angle, and chunks were attached to the grid by redeposition welding at 0.3 nA with three multi-pass CCS patterns (3 × 0.8 × 5 µm) on either side of each chunk. The sample was then coated again by 60 s sputtered platinum, 60 s GIS organometallic platinum, and 60 s sputtered platinum.

Grids with deposited chunks were transferred to an Arctis PFIB/SEM (Thermo Fisher Scientific) for lamella preparation. *C. elegans* positions within each chunk were confirmed using the integrated fluorescence microscope (iFLM) module (reflection: 1% laser, 5ms exposure; GFP: 10% laser, 700 ms exposure; both at binning 2). Chunks were thinned to approximately 550 nm with AutoTEM Cryo 2.4 (Thermo Fisher Scientific) using xenon plasma with currents of 4 nA, 1 nA and 0.3 nA. Manual polishing was then performed at 0.1 nA and 30 pA.

### *C. elegans* tilt-series acquisition

Tilt-series were acquired on a Titan Krios G4 transmission electron microscope (Thermo Fisher Scientific) equipped with a Selectris X energy filter. Data were recorded on a Falcon 4i camera in EER format using electron counting mode. Data collection was performed in SerialEM (v4.2.1) using SPACEtomo (v1.3) (Eisenstein et al., 2024) with beam tilt compensation (using SerialEM’s Coma vs Image Shift calibration). Tilt series were collected using a dose-symmetric scheme with a tilt range of ±54°with 3°increments, using a starting tilt to correct for the lamella milling angle.

High-magnification tilt series were acquired a nominal magnification of 81,000x (calibrated pixel size 1.48 Å/px) using the SerialEM Low Dose Record preset. The electron dose was 4 e^—^/Å^2^/tilt image, with a target defocus range of -3 to -5 µm applied in 0.5 µm increments. Low-magnification tilt series were acquired using the SerialEM Low Dose Search preset at magnifications of 11,500! or 15,000! (pixel sizes 10.95 and 8.42 Å/px, respectively), with an electron dose of 0.2–0.3 e^—^/Å^2^/tilt image and a defocus offset of -20 to -30 µm relative to the high-magnification setting. Detailed acquisition parameters for specific datasets are summarised in Supplementary Table S2.

### *C. elegans* cryo-ET data processing

Gain correction, motion correction and CTF estimation of tilt images was performed using the WarpTools pipeline (Tegunov and Cramer, 2019). Tilt images in which grid bars entered the field of view were manually excluded using custom scripts. Tilt-series alignment and tomogram reconstruction was performed using Are-Tomo (v1.2.5) (Zheng et al., 2022) with binning factors of 2 or 4. Tomograms were then flipped using IMOD (Mastronarde and Held, 2017) to obtain the correct map handedness and low-pass filtered. Prior to segmentation, tomograms were denoised with either CryoCARE (Buchholz et al., 2018) or IsoNet2 (Liu et al., 2025).

Membranes were segmented using MemBrain (Lamm et al., 2022). Lamella boundaries were annotated on the original tomograms in IMOD (Kremer et al., 1996) on three XZ slices, after which custom scripts were used to interpolate between these annotations and generate a binary lamella mask. This mask was then applied to the segmented volumes to mask to the lamella region. Membrane segmentations were separated by connected components and subsequently grouped according to their biological identity. Additional cellular features (muscle filaments, microtubules, ribosomes, ATPases, granules) were segmented using Dragonfly (Object Research Systems, v2025.1). For this, manual segmentation of patches from four slices of each tomogram was used to train a model via the Segmentation Wizard, which was then applied to predict segmentations across the full tomogram. The binary lamella masks generated for membrane segmentation were also applied to the Dragonfly-derived segmentations. Regions corresponding to ice contamination or GIS layer artifacts were manually removed using the Volume Eraser tool in ChimeraX. Final 3D segmentations were visualised in ChimeraX (Pettersen et al., 2021).

## ACKNOWLEDGEMENTS

We would like to thank Dr Susanna Cogo and Dr Eva Kevei from the University of Reading for providing *C. elegans* strains. We would also like to thank Dr Matthew Case and Dr Thomas Glen for microscope support. This work was supported by the Wellcome Trust through the Electrifying Life Science project (220526/Z/20/Z), Innovate UK Horizon Europe Guarantee “IMAGINE” (10048356) and a Wellcome Career Development Award (225902/Z/22/Z to M.G.). The Rosalind Franklin Institute is funded by UK Research and Innovation through the Engineering and Physical Sciences Research Council (EPSRC).

## AUTHOR CONTRIBUTIONS

**H.W**.: Conceptualisation; software development (initial scripts implementing interleaved multi-magnification acquisition within PACEtomo); sample preparation (*E. coli*); lamella preparation (*E. coli* and *C. elegans*); cryo-electron tomography data collection; data analysis; writing (original draft, editing and review). **V.G.G**.: Sample preparation (*C. elegans*); serial lift-out of *C. elegans*; lamella preparation (*C. elegans*). **F.E**.: Software development (implementation of multi-magnification acquisition into SPACEtomo). **M.G**.: Writing (editing and review); supervision; funding. All authors reviewed the manuscript.

## CODE AVAILABILITY

The SerialEM Python scripts used in this study are publicly available on GitHub. Target selection and setup were performed using SPACEtomo (https://github.com/eisfabian/SPACEtomo), and tilt-series acquisition was carried out using PACEtomo (https://github.com/eisfabian/PACEtomo).

## DATA AVAILABILITY

Subtomogram averages were deposited in the Electron Microscopy Data Bank (EMDB) under the accession codes EMD-XXXXX (interleaved) and EMD-XXXXX (non-interleaved). Raw microscopy data and associated metadata were deposited in the Electron Microscopy Public Image Archive (EMPIAR) under the accession code EMPIAR-XXXXX.

## References

Berger, C., Watson, H., Naismith, J. H., Dumoux, M., and Grange, M. (2025). Xenon plasma focused ion beam lamella fabrication on high-pressure frozen specimens for structural cell biology. Nature Communications 2025 16:1, 16:1–12. doi: 10.1038/s41467-025-57493-3.

Bouvette, J., Liu, H. F., Du, X., Zhou, Y., Sikkema, A. P., da Fonseca Rezende e Mello, J., Klemm, B. P., Huang, R., Schaaper, R. M., Borgnia, M. J., and Bartesaghi, A. (2021). Beam image-shift accelerated data acquisition for near-atomic resolution single-particle cryo-electron tomography. Nature Communications, 12(1):1–11. doi: 10.1038/S41467-021-22251-8.

Buchholz, T. O., Jordan, M., Pigino, G., and Jug, F. (2018). Cryo-CARE: Content-Aware Image Restoration for Cryo-Transmission Electron Microscopy Data. Proceedings - International Symposium on Biomedical Imaging, 2019-April:502–506. doi: 10.1109/ISBI.2019.8759519.

Chen, W. and Kudryashev, M. (2020). Structure of RyR1 in native membranes. The EMBO Reports 2020 21:5, 21(5):EMBR201949891–. doi: 10.15252/EMBR.201949891.

Chua, E. Y., Alink, L. M., Kopylov, M., Johnston, J. D., Eisenstein, F., and de Marco, A. (2024). Square beams for optimal tiling in transmission electron microscopy. Nature Methods 2024 21:4, 21(4):562–565. doi: 10.1038/s41592-023-02161-x.

Dickerson, J. L., Lu, P. H., Hristov, D., Dunin-Borkowski, R. E., and Russo, C. J. (2022). Imaging biological macromolecules in thick specimens: The role of inelastic scattering in cryoEM. Ultramicroscopy, 237:113510. doi: 10.1016/J.ULTRAMIC.2022.113510.

Downes, K., Flood, J., Nans, A., Verren, S. V. d., Audhya, A., and Zanetti, G. (2025). Multi-scale Molecular Imaging of Human Cells reveals COPI and COPII Vesicles at ER Exit Sites. bioRxiv, page 2025.07.29.667472. doi: 10.1101/2025.07.29.667472.

Du, D. X. and Fitzpatrick, A. W. (2023). Design of an ultrafast pulsed ponderomotive phase plate for cryo-electron tomography. Cell Reports Methods, 3(1):100387. doi: 10.1016/J.CRMETH.2022.100387.

Eisenstein, F., Yanagisawa, H., Kashihara, H., Kikkawa, M., Tsukita, S., and Danev, R. (2022). Parallel cryo electron tomography on in situ lamellae. Nature Methods 2022 20:1, 20: 131–138. doi: 10.1038/s41592-022-01690-1.

Eisenstein, F., Fukuda, Y., and Danev, R. (2024). Smart parallel automated cryo-electron tomography. Nature Methods 2024 21:9, 21:1612–1615. doi: 10.1038/s41592-024-02373-9.

Friedman, J. R. and Voeltz, G. K. (2011). The ER in 3D: a multifunctional dynamic membrane network. Trends in Cell Biology, 21(12):709–717. doi: 10.1016/J.TCB.2011.07.004.

Glynn, C., Smith, J. L., Case, M., Csöndör, R., Katsini, A., Sanita, M. E., Glen, T. S., Pennington, A., and Grange, M. (2025). A generalizable and targeted molecular biopsy approach for in situ cryogenic electron tomography of vitreous brain tissue. Cell Reports Methods, 5. doi: 10.1016/J.CRMETH.2025.101080/.

Grant, T. and Grigorieff, N. (2015). Measuring the optimal exposure for single particle cryo-em using a 2.6 Å reconstruction of rotavirus vp6. eLife, 4. doi: 10.7554/ELIFE.06980.

Hagen, W. J., Wan, W., and Briggs, J. A. (2017). Implementation of a cryo-electron tomography tilt-scheme optimized for high resolution subtomogram averaging. Journal of Structural Biology, 197:191–198. doi: 10.1016/J.JSB.2016.06.007.

Hylton, R., Sanders, M. B., Prajica, A., Rice, G., and Raunser, S. (2025). Streamlined montage cryo-electron tomography for exploring the ultrastructure of cells and tissues. bioRxiv, page 2025.09.01.673430. doi: 10.1101/2025.09.01.673430.

Kremer, J. R., Mastronarde, D. N., and McIntosh, J. R. (1996). Computer Visualization of Three-Dimensional Image Data Using IMOD. Journal of Structural Biology, 116(1): 71–76. doi: 10.1006/JSBI.1996.0013.

Lamm, L., Righetto, R. D., Wietrzynski, W., Pöge, M., Martinez-Sanchez, A., Peng, T., and Engel, B. D. (2022). MemBrain: A deep learning-aided pipeline for detection of membrane proteins in Cryo-electron tomograms. Computer Methods and Programs in Biomedicine, 224:106990. doi: 10.1016/J.CMPB.2022.106990.

Lange, F., Ratz, M., Dohrke, J. N., Le Vasseur, M., Wenzel, D., Ilgen, P., Riedel, D., and Jakobs, S. (2025). In situ architecture of the human prohibitin complex. Nature Cell Biology 2025 27:4, 27(4):633–640. doi: 10.1038/s41556-025-01620-1.

Leapman, R. D. and Sun, S. (1995). Cryo-electron energy loss spectroscopy: observations on vitrified hydrated specimens and radiation damage. Ultramicroscopy, 59:71–79. doi: 10.1016/0304-3991(95)00019-W.

Liu, Y. T., Zhang, H., Wang, H., Tao, C. L., Bi, G. Q., and Zhou, Z. H. (2022). Isotropic reconstruction for electron tomography with deep learning. Nature Communications 2022 13:1, 13(1):1–17. doi: 10.1038/s41467-022-33957-8.

Liu, Y.-T., Fan, H., Jih, J., Tran, L., Zhang, X., and Zhou, Z. H. (2025). IsoNet2 determines cellular structures at submolecular resolution without averaging. bioRxiv, page 2025.12.09.693325. doi: 10.64898/2025.12.09.693325.

Mastronarde, D. N. (2005). Automated electron microscope tomography using robust prediction of specimen movements. Journal of Structural Biology, 152:36–51. doi: 10.1016/J.JSB.2005.07.007.

Mastronarde, D. N. and Held, S. R. (2017). Automated tilt series alignment and tomographic reconstruction in IMOD. Journal of Structural Biology, 197(2):102–113. doi: 10.1016/J.JSB.2016.07.011.

Nguyen, H. T. D., Perone, G., Klena, N., Vazzana, R., Don, F. K., Silva, M., Sorrentino, S., Swuec, P., Leroux, F., Kalebic, N., Coscia, F., and Erdmann, P. S. (2024). Serialized ongrid lift-in sectioning for tomography (solist) enables a biopsy at the nanoscale. Nature Methods, 21:1693–1701. doi: 10.1038/S41592-024-02384-6.

Nikolay, R., Hilal, T., Qin, B., Mielke, T., Bürger, J., Loerke, J., Textoris-Taube, K., Nierhaus, K. H., and Spahn, C. M. (2018). Structural Visualization of the Formation and Activation of the 50S Ribosomal Subunit during In Vitro Reconstitution. Molecular Cell, 70(5):881–893. doi: 10.1016/j.molcel.2018.05.003.

Peck, A., Carter, S. D., Mai, H., Chen, S., Burt, A., and Jensen, G. J. (2022). Montage electron tomography of vitrified specimens. Journal of Structural Biology, 214:107860. doi: 10.1016/J.JSB.2022.107860.

Petrov, P. N., Zhang, J. T., Axelrod, J. J., and Müller, H. (2024). Crossed laser phase plates for transmission electron microscopy. arXiv.

Pettersen, E. F., Goddard, T. D., Huang, C. C., Meng, E. C., Couch, G. S., Croll, T. I., Morris, J. H., and Ferrin, T. E. (2021). UCSF ChimeraX: Structure visualization for researchers, educators, and developers. Protein Science, 30(1):70–82. doi: 10.1002/PRO.3943.

Puhka, M., Joensuu, M., Vihinen, H., Belevich, I., and Jokitalo, E. (2012). Progressive sheet-to-tubule transformation is a general mechanism for endoplasmic reticulum partitioning in dividing mammalian cells. Molecular Biology of the Cell, 23(13):2424–2432. doi: 10.1091/MBC.E10-12-0950.

Qadota, H., Matsunaga, Y., Nguyen, K. C., Mattheyses, A., Hall, D. H., and Benian, G. M. (2017). High-resolution imaging of muscle attachment structures in Caenorhabditis elegans. Cytoskeleton, 74(11):426–442. doi: 10.1002/CM.21410;.

Rosenthal, P. B. and Henderson, R. (2003). Optimal Determination of Particle Orientation, Absolute Hand, and Contrast Loss in Single-particle Electron Cryomicroscopy. Journal of Molecular Biology, 333(4):721–745. doi: 10.1016/J.JMB.2003.07.013.

Schiøtz, O. H., Kaiser, C. J., Klumpe, S., Morado, D. R., Poege, M., Schneider, J., Beck, F., Klebl, D. P., Thompson, C., and Plitzko, J. M. (2024). Serial lift-out: sampling the molecular anatomy of whole organisms. Nature Methods, 21:1684–1692. doi: 10.1038/S41592-023-02113-5.

Schorb, M., Haberbosch, I., Hagen, W. J., Schwab, Y., and Mastronarde, D. N. (2019). Software tools for automated transmission electron microscopy. Nature Methods, 16: 471–477. doi: 10.1038/S41592-019-0396-9.

Schwartz, O., Axelrod, J. J., Campbell, S. L., Turnbaugh, C., Glaeser, R. M., and Müller, H. (2019). Laser phase plate for transmission electron microscopy. Nature Methods, 16 (10):1016–1020. doi: 10.1038/S41592-019-0552-2.

Shemesh, T., Klemm, R. W., Romano, F. B., Wang, S., Vaughan, J., Zhuang, X., Tukachinsky, H., Kozlov, M. M., and Rapoport, T. A. (2014). A model for the generation and interconversion of ER morphologies. Proceedings of the National Academy of Sciences of the United States of America, 111(49):E5243–E5251. doi: 10.1073/PNAS.1419997111.

Strauss, M., Hofhaus, G., Schröder, R. R., and Kühlbrandt, W. (2008). Dimer ribbons of ATP synthase shape the inner mitochondrial membrane. EMBO Journal, 27(7):1154–1160. doi: 10.1038/EMBOJ.2008.35.

Tegunov, D. and Cramer, P. (2019). Real-time cryo-electron microscopy data preprocessing with Warp. Nature Methods 2019 16:11, 16(11):1146–1152. doi: 10.1038/s41592-019-0580-y.

Tegunov, D., Xue, L., Dienemann, C., Cramer, P., and Mahamid, J. (2021). Multi-particle cryo-EM refinement with M visualizes ribosome-antibiotic complex at 3.5 Å in cells. Nature Methods, 18(2):186–193. doi: 10.1038/S41592-020-01054-7.

Wagner, T., Merino, F., Stabrin, M., Moriya, T., Antoni, C., Apelbaum, A., Hagel, P., Sitsel, O., Raisch, T., Prumbaum, D., Quentin, D., Roderer, D., Tacke, S., Siebolds, B., Schubert, E., Shaikh, T. R., Lill, P., Gatsogiannis, C., and Raunser, S. (2019). SPHIRE-crYOLO is a fast and accurate fully automated particle picker for cryo-EM. Communications Biology, 2(1):1–13. doi: 10.1038/S42003-019-0437-Z.

Wu, J., Liu, C., Wang, A. J., Gao, Y. Z., Fu, L. T., Liu, Z., Dickerson, J. L., Russo, C. J., and Wang, P. (2025). Chromatic aberration (Cc) corrected cryo-EM: The structure of pseudorabies virus (PRV) using both zero-loss and energy loss electrons. Ultramicroscopy, 276:114182. doi: 10.1016/J.ULTRAMIC.2025.114182.

Xing, H., Taniguchi, R., Khusainov, I., Kreysing, J. P., Welsch, S., Turoňová, B., and Beck, M. (2023). Translation dynamics in human cells visualized at high resolution reveal cancer drug action. Science, 381(6653):70–75. doi: 10.1126/SCIENCE.ADH1411.

Xue, L., Spahn, C. M., Schacherl, M., and Mahamid, J. (2025). Structural insights into context-dependent inhibitory mechanisms of chloramphenicol in cells. Nature Structural and Molecular Biology, 32(2):257–267. doi: 10.1038/S41594-024-01441-0.

Yang, J. E., Larson, M. R., Sibert, B. S., Kim, J. Y., Parrell, D., Sanchez, J. C., Pappas, V., Kumar, A., Cai, K., Thompson, K., and Wright, E. R. (2023). Correlative montage parallel array cryo-tomography for in situ structural cell biology. Nature Methods 2023 20:10, 20: 1537–1543. doi: 10.1038/s41592-023-01999-5.

Zheng, S., Wolff, G., Greenan, G., Chen, Z., Faas, F. G., Bárcena, M., Koster, A. J., Cheng, Y., and Agard, D. A. (2022). AreTomo: An integrated software package for automated marker-free, motion-corrected cryo-electron tomographic alignment and reconstruction. Journal of Structural Biology: X, 6:100068. doi: 10.1016/J.YJSBX.2022.100068.

