## Supplementary Information for "Interleaved multi-magnification cryo-electron tomography bridges cellular and structural biology"

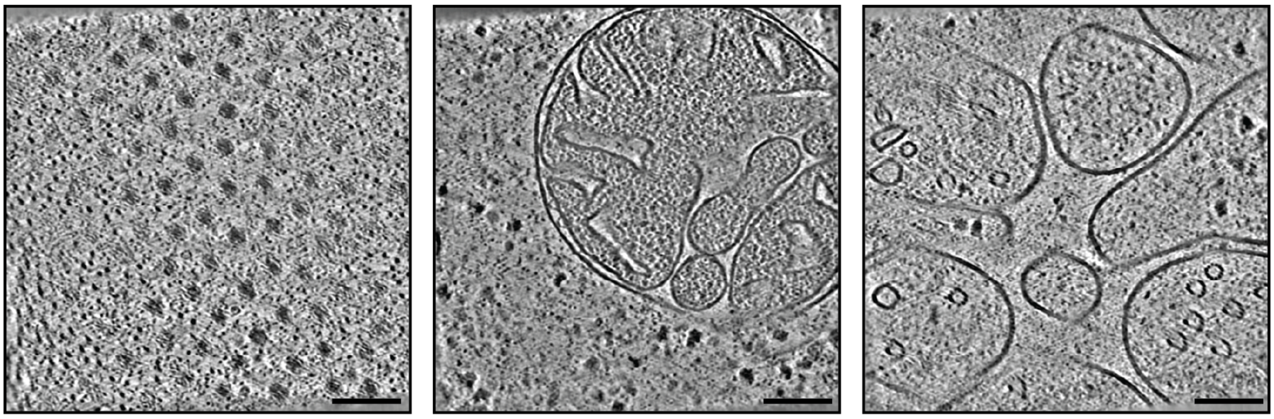

**Figure S1.** Additional denoised HM tomogram slices for the regions indicated in Figure 1B. Scale bar: 100 nm

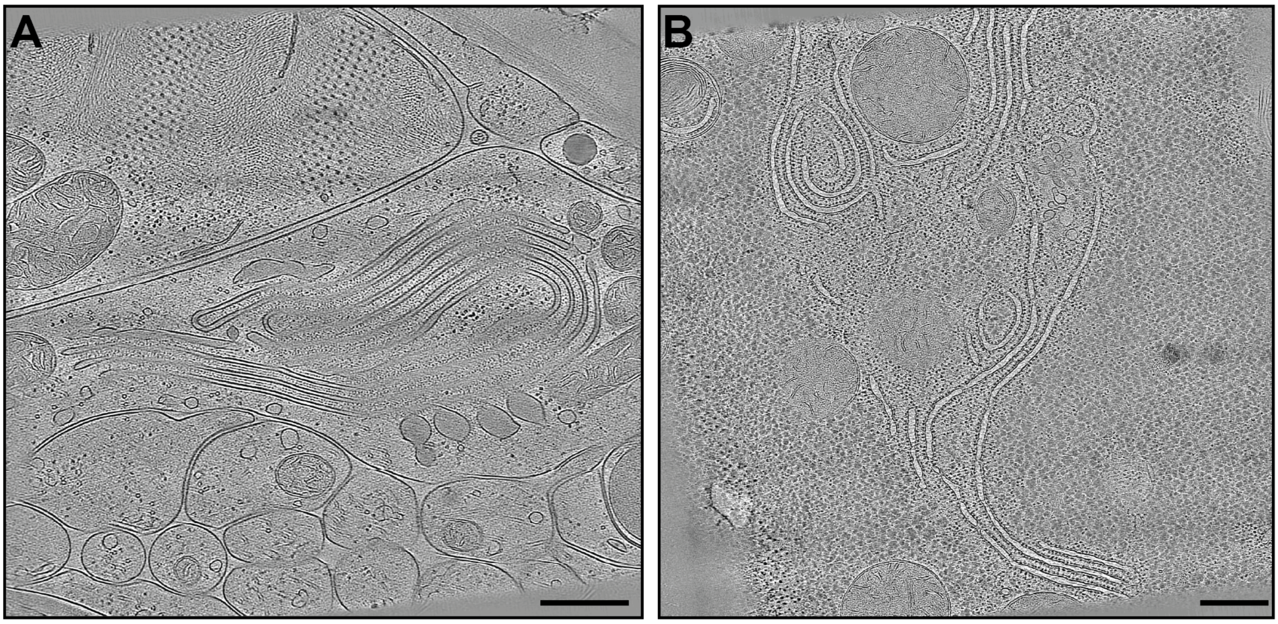


**Figure S2.** LM tomogram slices of *C. elegans*, corresponding to the segmentations in Figure 2A and Figure 2C respectively. Scale bars: 500 nm.

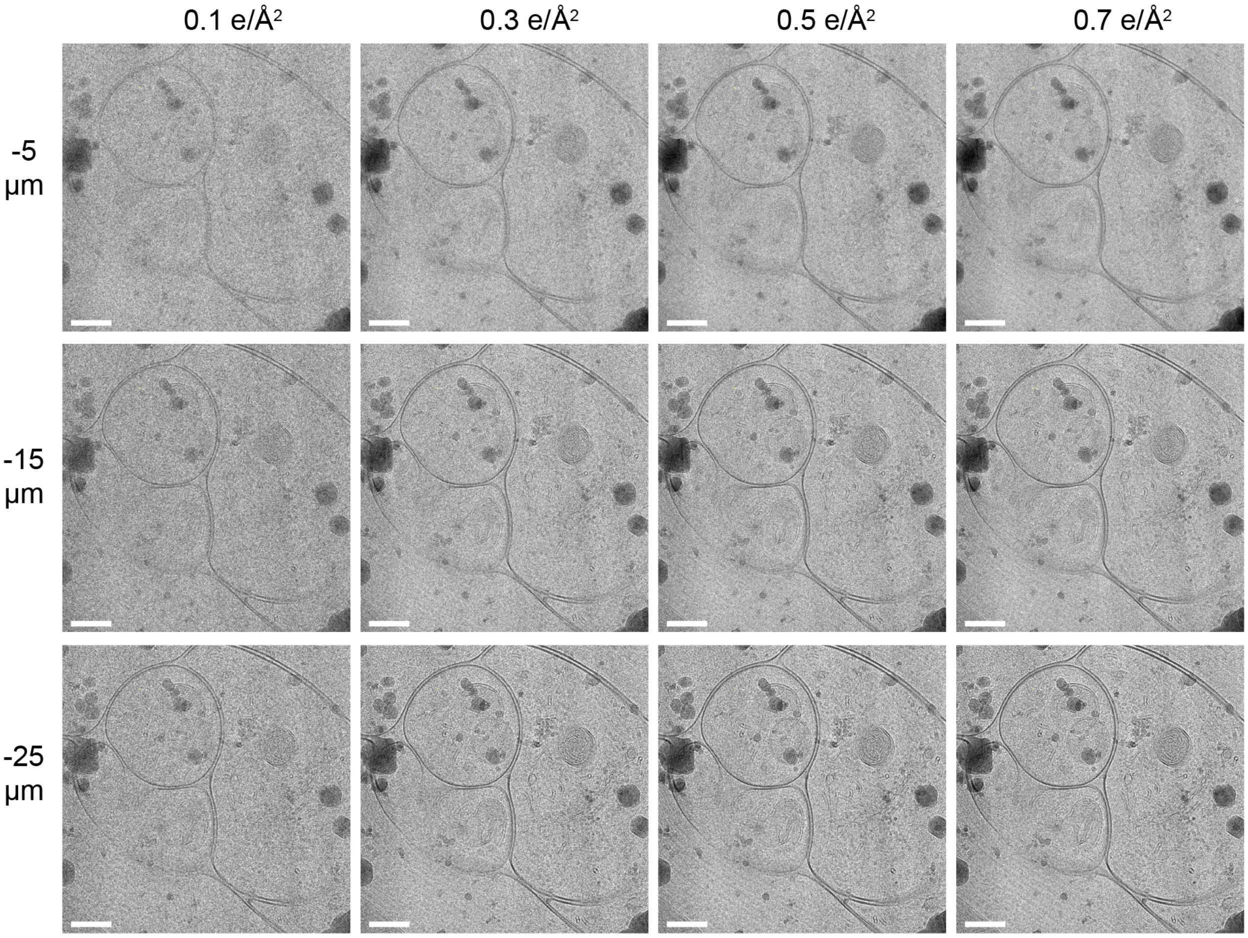


**Figure S3.** Comparison of TEM images at 11,500x magnification (11.0 Å/px, cropped FOV) at varied combinations of dose and defocus. Scale bars: 100 nm

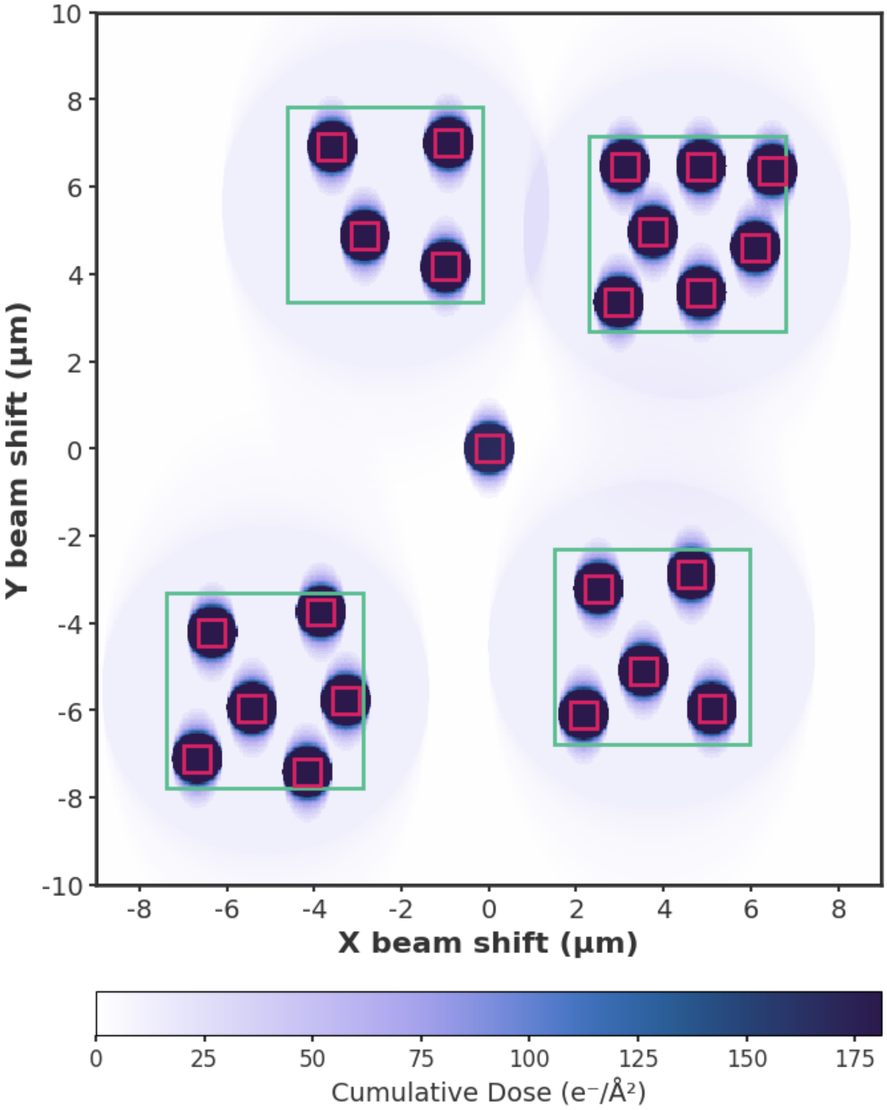


**Figure S4. Cumulative dose map of the lamella during tilt series acquisition.** The heatmap shows the spatial variation in accumulated dose ($e^-/Å^2$), calculated from all acquisition positions (positions as shown in Figure 3C) using dose rates of 4 $e^-/Å^2$/tilt for high magnification and 0.3 $e^-/Å^2$/tilt for low magnification. Squares indicate the detector field of view at 0° tilt for each magnification mode. Beam shift distances are relative to the central tracking/focus target area.

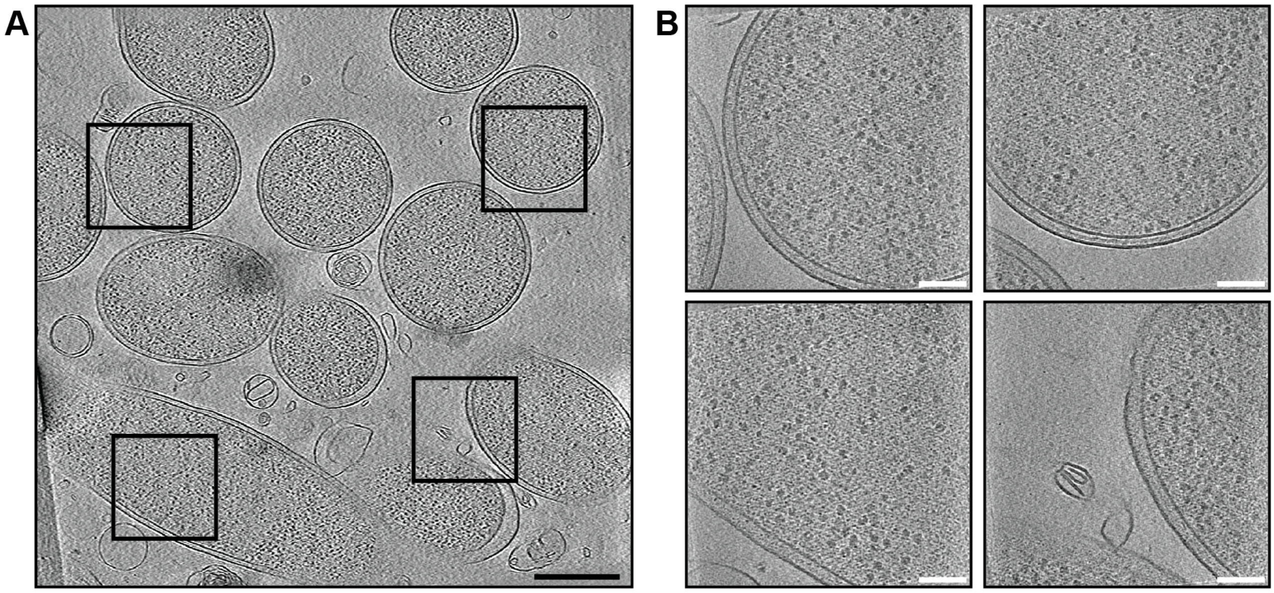

**Figure S5. Representative *E. coli* data.**
Representative *E. coli* data. Tomogram slices of **(A)** low-magnification and **(B)** high-magnification data. The positions of the HM tomograms are indicated with black boxes on (A). Scale bars: 500 nm and 100 nm, respectively.

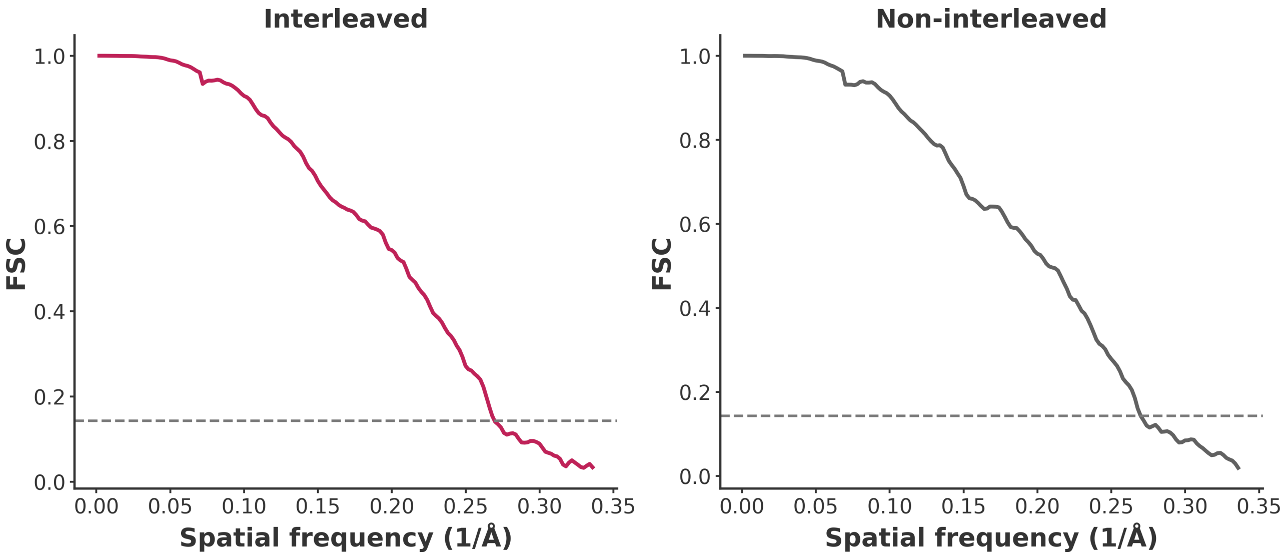


**Figure S6.** Randomised particle half-map FSC curves for interleaved and non-interleaved *E. coli* 50S reconstructions. The corrected FSC is shown as a function of spatial frequency (1/Å) with resolution estimated using the 0.143 criterion (dashed line).

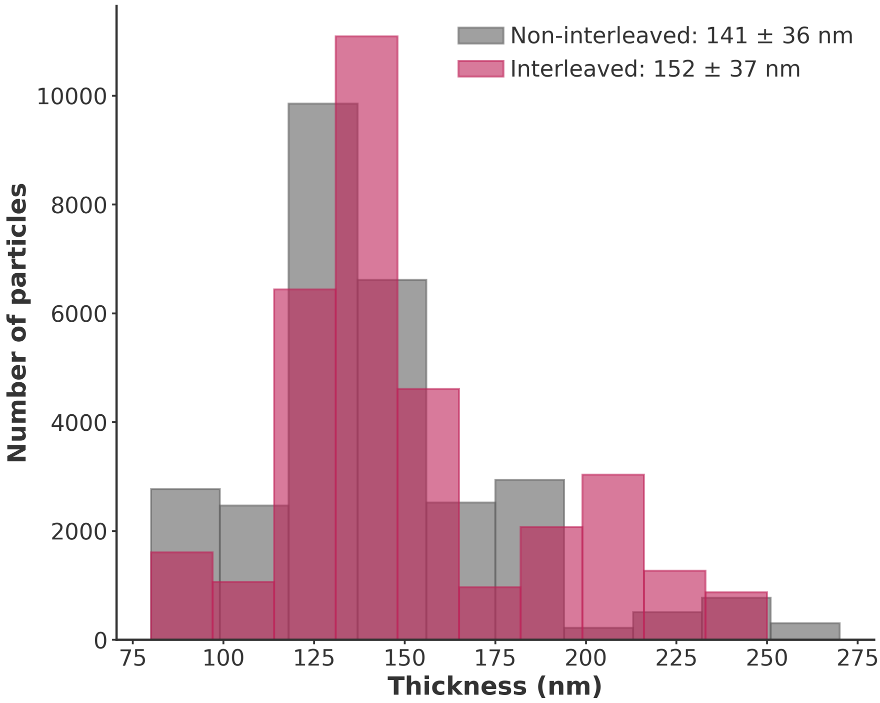


**Figure S7.** Distribution of tomogram thickness for interleaved and non-interleaved datasets, per particle, used for B-factor analysis.

### Supplementary Tables

**Table S1.** Summary of typical imaging parameters for high (HM) and low (LM) magnification tilt-series acquisition

| Parameter | High Magnification (HM) | Low Magnification (LM) |
|---|---|---|
| Nominal magnification | 64k – 105k | 8.7k – 19.5k |
| Pixel size (Å/px) | 1.9 - 1.2 Å/px | 14.5 – 6.3 Å/px |
| Field of view ($\mu m^2$) | 0.6 - 0.2 $\mu m^2$ | 35.3 - 6.7 $\mu m^2$ |
| Defocus range | -3 – -5 µm | > -15 µm |
| Dose per tilt image | 3 – 5 $e^-/Å^2$ | 0.2 – 0.5 $e^-/Å^2$ |
| Total dose per tilt series | 120 – 200 $e^-/Å^2$ | 8 – 20 $e^-/Å^2$ |

**Table S2.** Acquisition parameters for *E. coli* and *C. elegans*

| Parameter | ***E. coli*** | ***E. coli*** | ***C. elegans*** | ***C. elegans* (BWM)** | ***C. elegans* (RER)** |
|---|---|---|---|---|---|
| Nominal magnification | 81,000x | 15,000x | 81,000x | 15,000x | 11,500x |
| Pixel size (Å/px) | 1.48 | 8.42 | 1.48 | 8.42 | 10.95 |
| Energy filter slit width (eV) | 10 | 20 | 10 | 20 | 20 |
| Electron dose per tilt ($e/Å^2$) | 4 | 0.5 | 4 | 0.3 | 0.2 |
| Tilt range (°) | ±54 | ±54 | ±54 | ±54 | ±60 |
| Tilt increment (°) | 3 | 3 | 3 | 3 | 3 |
| Total electron dose ($e/Å^2$) | 148 | 18.5 | 148 | 11.1 | 8.2 |
| Defocus range (µm) | -3 to -5 | -23 to -25 | -3 to -5 | -33 to -35 | -23 to -25 |

**Table S3.** Subtomogram averaging statistics for *E. coli* ribosomes

| Parameter | Interleaved | Non-interleaved |
|---|---|---|
| Number of tilt-series | 98 | 83 |
| Number of particles | 33,035 | 28,981 |
| FSC 0.143 resolution (Å) | 3.7 | 3.7 |
| B-factor ($Å^2$) | 96 | 92 |
| EMDB | | |
